# Antagonistic pathogen-mediated selection favours the maintenance of innate immune gene polymorphism in a widespread wild ungulate

**DOI:** 10.1101/458216

**Authors:** Erwan Quéméré, Pauline Hessenauer, Maxime Galan, Marie Fernandez, Joël Merlet, Yannick Chaval, Nicolas Morellet, Hélène Verheyden, Emmanuelle Gilot-Fromont, Nathalie Charbonnel

**Affiliations:** Université de Toulouse, INRAE, CEFS, 31320, Castanet-Tolosan, France; LTSER ZA PYRénées GARonne, 31320, Auzeville-Tolosane, France; ESE, Ecology and Ecosystems Health, Agrocampus Ouest, INRAE, 35042 Rennes, France; Département de Foresterie, Université Laval, Quebec, Canada; CBGP, INRAE, CIRAD, IRD, Montpellier SupAgro, Univ Montpellier, Montpellier, France; Université de Lyon, Université Lyon 1, UMR CNRS 5558, Villeurbanne, France; Université de Lyon, VetAgro Sup, Marcy l’Etoile, France

**Keywords:** Toll-like genes, antagonistic effects, balancing selection, habitat heterogeneity, roe deer

## Abstract

Toll-like Receptors (TLR) play a central role in recognition and host frontline defence against a wide range of pathogens. A number of recent studies have shown that TLR genes (*Tlrs*) often exhibit a large polymorphism in natural populations. Yet, there is little knowledge on how this polymorphism is maintained and how it influences disease susceptibility in the wild. In a previous work, we showed that some *Tlrs* exhibit similarly high levels of genetic diversity than *Mhc* and contemporary signatures of balancing selection in roe deer (*Capreolus capreolus*), an abundant and widespread ungulate in Europe. *Here, we tested whether Mhc-Drb* or *Tlr* (*Tlr2, Tlr4* and *Tlr5*) diversity is driven by *pathogen-mediated selection. We* examined the relationships between their genotype (heterozygosity status and presence of specific alleles) and infections with *Toxoplasma* and *Chlamydia*, two intracellular pathogens known to cause reproductive failure in ungulates. We showed that *Toxoplasma* and *Chlamydia* exposures vary significantly across year and landscape structure with few co-infection events detected, and that the two pathogens act antagonistically on *Tlr2* polymorphism. By contrast, we found no evidence of association with *Mhc-Drb* and a limited support for *Tlr* heterozygosity advantage. Our study confirmed the importance of looking beyond *Mhc* genes in wildlife immunogenetic studies. It also emphasized the necessity to consider multiple pathogen challenges and their spatiotemporal variation to improve our understanding of vertebrate defence evolution against pathogens.

## 1. INTRODUCTION

While evolutionary theory predicts that selection and random drift should deplete genetic diversity, a number of genes and genomic regions have maintained abundant polymorphism in natural populations with exceptionally high ratio of non-synonymous (protein-altering) to synonymous substitutions (Trowsdale and Parham, 2004; Hedrick, 2006). Balancing selection is a collective term for evolutionary processes that adaptively maintain variation in populations. It is a widespread process driving the diversity of genes involved in numerous biological functions including pathogen resistance (Sommer, 2005), color polymorphism (Wellenreuther, 2017) or mate choice/male competiveness (Moore and Moore, 1999; Johnston *et al*., 2013). There is an intense debate about the mechanisms driving balancing selection, with three primary hypotheses (putting aside “sexual selection”): the “heterozygote advantage” (Lewontin and Hubby, 1966; Doherty and Zinkernagel, 1975), the “fluctuating selection” over time and space (Levene, 1953; Hill, 1991) and the “negative frequency-dependent selection” (also called rare-allele advantage) (Takahata and Nei, 1990; Phillips *et al*., 2018). Disentangling these three mechanisms in natural systems is particularly challenging since they can operate in synergy and are not-mutually exclusive (Spurgin and Richardson, 2010).

Immunity-related genes are ideal candidates for studying how adaptive genetic diversity is maintained in wild populations because of their direct effect on survival (Møller and Saino, 2004) or reproductive success (Pedersen and Greives, 2008). The genes of the major histocompatibility complex (MHC) are often used as a model since they show an exceptional polymorphism in vertebrates (Spurgin and Richardson, 2010). Their structure and mechanisms of interaction with antigens are well described (Klein & Figueroa, 1986). Furthermore, their ability to trigger a rapid adaptive response to varying pathogen-mediated selective pressures (PMS) has been demonstrated experimentally (Eizaguirre *et al*., 2012; Phillips *et al*., 2018). Over the last few years, a growing body of literature identified PMS at *Mhc* genes in a large range of taxa (see Spurgin & Richardson, 2010 for a review) but few of them convincingly elucidated the relative influence of the three mechanisms mentioned above (but see Oliver et al., 2009; Philips et al., 2018). One likely reason is that most studies analysed associations between a single gene (most often a *Mhc* gene) and only one pathogen. However, decreasing sequencing costs now offer new opportunities to examine multiple genes and diseases in a single study, leading to a more realistic view of the functioning of the immune system and of the ecological context in which it evolves.

*Mhc*, although important, only corresponds to a fraction of the genetic variation in pathogen resistance (Acevedo-Whitehouse and Cunningham, 2006). Before the adaptive immunity has an opportunity to intervene in the host response, invading pathogens interact with different effectors of innate immunity that recognize specific structures (Jepson *et al*., 1997; Lam-Yuk-Tseung and Gros, 2003). Therefore many genes encoding proteins involved in pathogen recognition or immune regulation may be associated with the resistance against the same pathogen (Turner *et al*., 2011). In particular, genes encoding Toll-like receptors (TLR) play a key role in the host frontline defence against a wide range of microparasites such as fungi, bacteria, protozoa. This family of genes conserved in both invertebrate and vertebrate lineages are critical for the stimulation of many immune pathways such as inflammation, innate or acquired immunity (Akira *et al*., 2001). A multitude of studies have revealed associations between *Tlr* gene polymorphism and infectious diseases in humans and domestic species (Mun *et al*., 2003; Yang and Joyee, 2008). However, very few (most focusing on birds or rodents) have investigated their role in host-pathogen interactions in natural environment and their contemporary mechanisms of evolution at a population level (but see Kloch et al., 2018; Tschirren et al., 2013).

Moreover, infection with multiple pathogens is a real world rule (Maizels and Nussey, 2013) and some selective mechanisms such as heterozygous advantage or fluctuating selection may better be detected when accounting for the multi-pathogen context (Wegner *et al*., 2003; Sin *et al*., 2014). Most immunity-related genes have pleiotropic effects on resistance against several pathogens (Hedrick, 1999) and the maintenance of ‘susceptible’ (deleterious) alleles in a wild population can be due to antagonist effects of a given immunity-related gene on several pathogens (*i*.*e*. antagonistic pleiotropy). To date, only few studies have examined the selective effects of multiple pathogens on host immunogenetic patterns and the great majority focused on *Mhc* genes (Tollenaere et al., 2008 on voles; Loiseau et al., 2008 on house sparrows; Kamath et al., 2014 in plain zebras; Froeschke & Sommer, 2012 on striped mouse but see Dubois *et al*., 2017, Antonides et al., 2019).

Here, we investigated the selective mechanisms by which multiple pathogens could drive the genetic diversity of immunity-related genes in a free-ranging population of European roe deer (*Capreolus capreolus*) inhabiting an agricultural landscape in Southwestern France. We focused on four genes encoding pattern-recognition receptors (PRRs) that detect specific microbes or pathogen-associated molecular patterns: the widely-used *Mhc class II-Drb* gene and three non-*Mhc* genes encoding Toll-like receptors genes (*Tlr2, Tlr4, Tlr5*). In a recent work, we showed that these genes exhibit high levels of amino acid diversity within French roe deer populations including the population studied here, hence suggesting an important functional repertoire (Quéméré *et al*., 2015). In particular, *Tlr2* and *Tlr4* exhibited several haplotypes at moderate frequency and very low levels of differentiation among roe deer populations compared to the patterns obtained with neutral markers. These findings support the hypothesis that the polymorphism of these genes is shaped by balancing selection although the underlying mechanism(s) remains to be investigated.

The study population is heavily exposed to a range of pathogens that also infect livestock, including the abortive *Toxoplasma gondii* or *Chlamydia abortus* (Candela *et al*., 2014). These two pathogens have negative influence on the survival and reproductive performances of ungulates (Pioz *et al*., 2008; Dubey, 2009). It can therefore be assumed that they exert selective pressures on immunity-related genes and in particular on *Tlr* genes that play a major role in host defence against protozoans and gram-negative bacteria (Gazzinelli and Denkers, 2006; Yarovinsky, 2008; Miller *et al*., 2009; Beckett *et al*., 2012; Netea *et al*., 2012). To test this hypothesis, we first established whether there are biological (age, sex) and/or environmental patterns (year, landscape structure) in *Toxoplasma* and *Chlamydia* infectious status across hosts. Then, we analysed whether pathogen prevalence was associated with specific patterns of genetic variation observed at the four immunity-related genes, while accounting for biological/environmental heterogeneity. Specifically, we tested whether (i) particular alleles are associated with lower or higher pathogen prevalence (directional selection) and (i) heterozygous individuals exhibit a lower prevalence than homozygotes (heterozygote advantage). Support for the “heterozygote advantage” hypothesis has been found in many *Mhc* gene studies of mammals (Oliver *et al*., 2009; Hedrick, 2012; Sin *et al*., 2014), including wild ungulates (Paterson *et al*., 1998; Brambilla *et al*., 2018). The importance of this hypothesis for innate immunity-related genes has rarely been assessed (but see Antonides et al., 2019; Grueber et al., 2013) but a recent study in birds (Grueber, Wallis, & Jamieson, 2013) has found evidence for survival advantage associated with *Tlr4* heterozygote genotypes. Moreover, *Mhc* and *Tlr* genes are known to have pleiotropic effects on the resistance/susceptibility to pathogens harboring similar pathogen-associated molecular patterns (Hedrick, 1999; Kaisho and Akira, 2004; Loiseau *et al*., 2008). Since *Toxoplasma* and *Chlamydia* employ similar strategies to invade host cells (Romano and Coppens, 2013), we expect that they should both exert selective pressures on same genes, but with potentially antagonistic effects if the same allele confer resistance to a pathogen and susceptibility to the other (i.e. “antagonistic pleiotropy”, Kubinak *et al*., 2012).

## 2. MATERIAL AND METHODS

### 2.1 Study population and data collection

The study focused on a roe deer population inhabiting a heterogeneous agricultural landscape in Southern France (43°13⍰N, 0°52⍰E). This area called “Vallons et Côteaux de Gascogne” (VCG) is part of the Atelier Pyrénées Garonne (https://pygar.omp.eu/). Individuals were caught by drive-netting during winter (from January to March) between 2008 and 2016. The sampling area consisted of three sectors with contrasting landscape structure with regard to the proportion of woodland: a “closed” sector with two forest blocks, an “open” landscape including mainly meadows, crops and pastures with few fragmented woodlots, and an “intermediate” sector with inter-connected woodland fragments (Hewison *et al*., 2009). Deer from these three sectors belong to the same panmictic population (Coulon *et al*., 2006; Gervais *et al*., 2019). Individual sex, age class (juveniles: ≤ 1 year of age versus adults: > 1 year of age) and body mass were recorded. Blood samples were collected for pathogen screening (see below) and ear punches for genetic analyses. In total, we gathered samples from 433 annual captures corresponding to 328 different individuals (190 females and 138 males, 164 juveniles and 164 adults) among which 157 deer had been included in a previous immunogenetic studies (Quéméré et al., 2015). All applicable institutional and European guidelines for the care and use of animals were followed. The study was permitted by the land manager. All the procedures involving animals were approved by the Ethical Committee 115 of Toulouse and authorized by the French Ministry in charge of ethical evaluation (n° APAFIS#7880-2016120209523619v5).

### 2.2 *Tlr* and *Mhc-Drb* genotyping

The *Mhc-Drb* class II and *Tlr* (*Tlr2, Tlr4* and *Tlr5*) genes of 157 roe deer from VCG had been genotyped in a previous study (Quéméré *et al*., 2015). We completed this dataset by genotyping 171 new individuals using exactly the same procedure. DNA was extracted from alcohol-preserved tissues using the DNeasy Blood and Tissue kit (QIAGEN). *Tlr* genes were genotyped using a two-step approach: a pilot study on 32 individuals was first performed to identify polymorphic sites (SNPs) and linkage-disequilibrium (LD) groups. We screened almost the entire coding region of the three *Tlr* genes (82% in average) including the leucine-rich extracellular region of receptors involved in antigen-binding. Detailed on primer sequences, SNP positions and codon changes are provided in Quéméré et al. (2015). We then selected one SNP per LD group (primarily targeting non-synonymous sites) that was genotyped for all individuals using the KASPar allele-specific genotyping system provided by KBiosciences (Hoddesdon, UK). A total of 13 SNPs were typed including 5, 3 and 5 SNPs for *Tlr2, Tlr4* and *Tlr5* respectively. Details on SNP position and codon change can be found in Table S1. Lastly, haplotypes were reconstructed from the phased SNPs using the procedure implemented in DNASP v5 (Librado & Rozas, 2008). The second exon of the *Mhc-Drb* class II gene encoding the ligand-binding domain of the protein was amplified and sequenced using Illumina MiSeq system as detailed in Quéméré et al. (2015). Haplotypes and individual genotypes were identified using the SESAME barcode software (Piry *et al*., 2012). All sequences have been submitted to NCBI Genbank (Accession nos. are in Table S2, Supporting information).

### 2.3 Pathogen screening

The serological status of roe deer for *Toxoplasma gondii* and/or *Chlamydia abortus* was analysed using classical enzyme-linked immunosorbent assays (ELISA) with specific commercial kits (Sevila *et al*., 2014). The specificity and sensitivity of these kits were respectively 97.4 and 99.4 % for *T. gondii* and 92.2 and 89 % for *C. abortus*. Although the kits were developed for domestic ruminants, they were shown to be reliable and efficient in wild ungulates (Gotteland *et al*., 2014) with a good concordance with classical test (e.g. high concordance (0.8) between ELISA and the Modified Agglutination Test, a reference test for *Toxoplasma* in roe deer). According to the manufacturer’s instructions, blood samples with antibody recognition level (ARL) > 30% (respectively 40%) were considered positive for *T. gondii* (respectively *C. abortus*) (see Sevila et al., 2014 for further details). In total, we obtained the *Toxoplasma* and *Chlamydia* serological status of 277 (with 74 repeated measures) and 196 (36 repeated measures) roe deer respectively (caught between 2008 and 2016, Table S3). The individual repeatability (R) and its standard error (SE) for *Toxoplasma* and *Chlamydia* seroprevalence were calculated using the R package ‘rptR’ (Stoffel *et al*., 2017). The proportion of individuals seropositive for both *Toxoplasma* and *Chlamydia* was examined in the years where both pathogens were screened (between 2008 and 2013).

### 2.4 Statistical analyses

We analysed the influence of immunity-related gene variation, environmental and life-history factors on the seroprevalence of *T. gondii* or *C. abortus* (binary response variable – presence/absence) using generalized linear mixed-effect models (GLMM) with a binomial family (using a logit link function). Analyses were performed using the *glmer* function implemented in the ‘lme4’ package (Bates *et al*., 2012) and ‘MuMin’ v1.7.7 (Barton, 2009) in R version 3.3.3 (R Development Core Team, 2017). In the first step, we investigated the effects of non-genetic biological and environmental factors that possibly affect pathogen exposure. The starting model included roe deer sex (male *versus* female) and age-class (juvenile *versus* adult) because both behaviour and host susceptibility may vary between sexes and across an individual’s lifetime (Klein, 2000) and because infection persistence is expected to result in a positive age-prevalence relationship (Gotteland *et al*., 2014). We also included the capture year (9 levels) and sectors (3 levels) as fixed effects to account for the inter-annual and among-landscapes variation in environmental conditions that may affect pathogen survival and thus encounter probability (e.g. climate, food resources, population density). Roe deer classically inhabit a single home range of relatively small (mean 0.76 km^2^) across their entire adult life (Morellet *et al*., 2013). In this study, roe deer home range was generally part of a unique capture sector. In all models, we included individual identity as a random effect to account for correlation amongst different observations of the same individual. We conducted an initial exploration of our data to ascertain their distribution and identify potential outliers (Zuur *et al*., 2009). We used a model selection procedure using Akaike information criterion with a correction for small sample sizes (AICC) to determine which model best explained variation in pathogen infection (models with lowest AIC). We only examined models with ΔAIC< 2 relative to the best model. Significant variables were retained in a reduced model for use in step 2.

In the second step, we investigated the “heterozygote advantage hypothesis” by fitting a binary fixed effect (heterozygote *vs* homozygote) and the “allele advantage” hypothesis by considering associations between the infection status and the number of copies (0, 1 or 2) of the most or second most frequent haplotype of *Tlr2, Tlr4, Tlr5* and *MHC-Drb*. We used variance inflation factors (VIF) to measure collinearity among predictors and retained variables with VIF values <3 (Zuur *et al*., 2009). To minimize multicollinearity, the effects of the two most frequent haplotypes of a gene and its heterozygosity were evaluated in separate models. The identity of year and sector (for *Toxoplasma*) and year (for *Chlamydia*) were included as random effects in all models because (i) individuals at a sector and/or year all experience similar conditions of exposure (see the results of step 1) and (ii) to account for spatial and/or temporal autocorrelation in allele frequencies that may lead to spurious associations with pathogen prevalence. In total, 256 candidate models were evaluated for both *Toxoplasma* and *Chlamydia* (see Table S4 for the full list of models).

## 3. RESULTS

### 3.1 Immunogenetic variation

Among the 293 tested individuals, we found nine functional alleles for *Mhc* class II-*Drb* gene, coding for different amino acid sequences. One haplotype (CacaDRB*0302) was common (60%), two other (CacaDRB*0201 and CacaDRB*0102) had intermediate frequency (18% and 10% respectively) while the six others were rare (<5%). Observed heterozygosity (*H*_*o*_) reached 0.55. We isolated six different haplotypes for *Tlr2* in 327 genotyped individuals, all corresponding to different amino acids sequences. Two haplotypes (*Tlr2-2*: “TGCCG” and *Tlr2-1*: “CATCG”) and showed particularly high frequency (51% and 35% respectively) while the four others were rare (< 5%). The two most common haplotypes belonged to two highly divergent genetic clusters and were separated by seven substitutions (including four non-synonymous ones) (Quéméré *et al*., 2015). The *Tlr4* gene exhibited four functional haplotypes in 327 individuals including two haplotypes (“CGG” and “GAG”) with moderate to high frequency (26% and 55% respectively) and two haplotypes with low frequencies (10% and 7%). Lastly, we isolated six haplotypes (in 324 individuals) for the *Tlr5* among which two (“GCTCG”, “ACCTG”) had high frequencies (38% and 31% respectively), one (“ATCCG”) had intermediate frequency (18%) and the three others were rare (<5%). Observed heterozygosity was relatively high for the three *Tlr* genes (*H*_*o*_ = 0.62, 0.64 and 0.69, *Tlr2, Tlr4* and *Tlr5* respectively).

### 3.2 Ecological patterns of parasitism

Seroprevalence of *Toxoplasma* reached 35% and was moderately repeatable among years at the individual level (R=0.21, SE=0.06 P-value=0.002) By comparison, seroprevalence of *Chlamydia* was lower (16%) but showed a higher individual consistency among capture events (R=0.70, SE=0.05, P-value<0.001). Among the 232 annual captures for which both pathogens were screened (196 different deer), only 18 individuals (7%) had anti-*Toxoplasma* and anti-*Chlamydia* antibodies simultaneously while 93 (40%) and 36 (15%) animals were seropositive solely for *Toxoplasma* and *Chlamydia* respectively (Table 1). We found a strong effect of “the year of capture” for the two pathogens with an increased probability of being seropositive for *Toxoplasma* in 2009 and 2010 (Seroprevalence = 83% and 56% respectively) and a decreased seroprevalence in 2009 for *Chlamydia* (16%) (Table 1). Additionally, we revealed a significant relationship between *Toxoplasma* infection status, age and landscape structure: antibody prevalence was higher in adults and in the “open” sector (Table S5, Table S6).

**Table 1.** Summary of the pathogen. data (number of samples per year in brackets). NA indicates the years for which no data were available.

| Pathogen | Number of samples | Number of individuals | Overall seroprevalence | Seroprevalence evaluated per year |  |  |  |  |  |  |  |  |
| --- | --- | --- | --- | --- | --- | --- | --- | --- | --- | --- | --- | --- |
|  |  |  |  | 2008 | 2009 | 2010 | 2011 | 2012 | 2013 | 2014 | 2015 | 2016 |
| *Toxoplasma* | 351 | 277 | 0.35 | 0.23<br>(40) | 0.83<br>(42) | 0.56<br>(43) | 0.24<br>(50) | 0.21<br>(42) | 0.24<br>(38) | 0.16<br>(31) | 0.3<br>(49) | 0.43<br>(16) |
| *Chlamydia* | 232 | 196 | 0.16 | 0.00<br>(40) | 0.14<br>(41) | 0.37<br>(43) | 0.28<br>(50) | 0.05<br>(42) | 0.00<br>(16) | NA | NA | NA |
| coinfection | 232 | 196 | 0.08 | 0<br>(40) | 5<br>(41) | 11<br>(43) | 2<br>(50) | 0<br>(42) | 0<br>(16) | NA | NA | NA |

### 3.3 Genetic effects on pathogen infection

The inclusion of *Tlr2* genotype significantly improved the *Toxoplasma* infection non-genetic model. The best-fitted model (ΔAICc = −5.71 with non-genetic model) (Table 2) included a positive effect of the number of copies of the “TGCCG” haplotype of *Tlr2* (*Tlr2-2*). Roe deer carrying two copies of this frequent haplotype (51%) showed a decreased *Toxoplasma* seroprevalence (odd ratio OR = 0.29 [0.11-0.77]) compared to individuals without this haplotype (Figure 1a, Table S7). Model including the second most-frequent haplotype “CATCG” (*Tlr2-1*) was poorly supported (ΔAICc = 3.58 with the best model): *Toxoplasma* prevalence substantially increased for deer carrying one copy of *Tlr2-1* (OR = 2.41 [1.15-5.07]) but this effect was not additive. Individuals with two copies showed similar seroprevalence (OR=2.21 [0.75-6.48]). In a post-hoc analysis, we re-run this model by including a dominant rather than an additive effect of *Tlr2-1* (presence/absence of the allele) and this greatly improved the model performance (ΔAICc = 1.56 with the best model) (Figure 1b). Deer carrying at least on Tlr2-1 copy had a decreased Toxoplasma seroprevalence. We did not detect any association between *Toxoplasma* infection and *Tlr4, Tlr5* or *Mhc-Drb* genetic variation.

**Table 2.** Performance of the best-fitted generalized linear mixed-effect models of *Toxoplasma* and *Chlamydia* seroprevalence. (with ΔAIC< 2 relative to the best model). ‘*Toxoplasma’* models all included *age* as a fixed factor and individual *ID, year* and *capture sectors* as random factors. ‘*Chlamydia’* models included animal ID and year as random factors. ‘n’ refers to the number of seroprevalence data. The selected models occur in bold. ‘k’ refers to the number of model parameters. ‘nb(*haplotype*)’ refers to the number of copies of the haplotype. h(*gene*) refers to the heterozygosity status of the gene (0/1).

| *Toxoplasma* (n=351) |  |  |  |  | *Chlamydia* (n=232) |  |  |  |  |
| --- | --- | --- | --- | --- | --- | --- | --- | --- | --- |
|  | k | AICc | ΔAICc | AICcWt |  | k | AICc | ΔAICc | AICcWt |
| nb(Tlr2-2) | 7 | 382.1 | -5.71 | 0.416 | nb(Tlr2-2)+nb(Tlr5-2) | 6 | 166.5 | -7.35 | 0.430 |
| nb(Tlr2-2)+h(Tlr4) | 8 | 383.6 | -4.28 | 0.204 | nb(Tlr2-1)+nb(Tlr5-2) | 6 | 167.8 | -5.96 | 0.215 |
| nb(Tlr2-2)+nb(Tlr4-4) | 9 | 383.7 | -4.15 | 0.192 | nb(Tlr2-2)+nb(Tlr5-2)+nb(Tlr4-4) | 8 | 168.1 | -5.71 | 0.190 |
| nb(Tlr2-2)+ h(Tlr5) | 8 | 383.7 | -4.11 | 0.188 | nb(Tlr2-2+nb(Tlr5-2)+h(Tlr4-4) | 7 | 168.4 | -5.44 | 0.165 |
| Non-genetic model | 5 | 387.65 | 0 |  | Non-genetic model | 2 | 173.7 | 0 |  |

**Figure 1.**
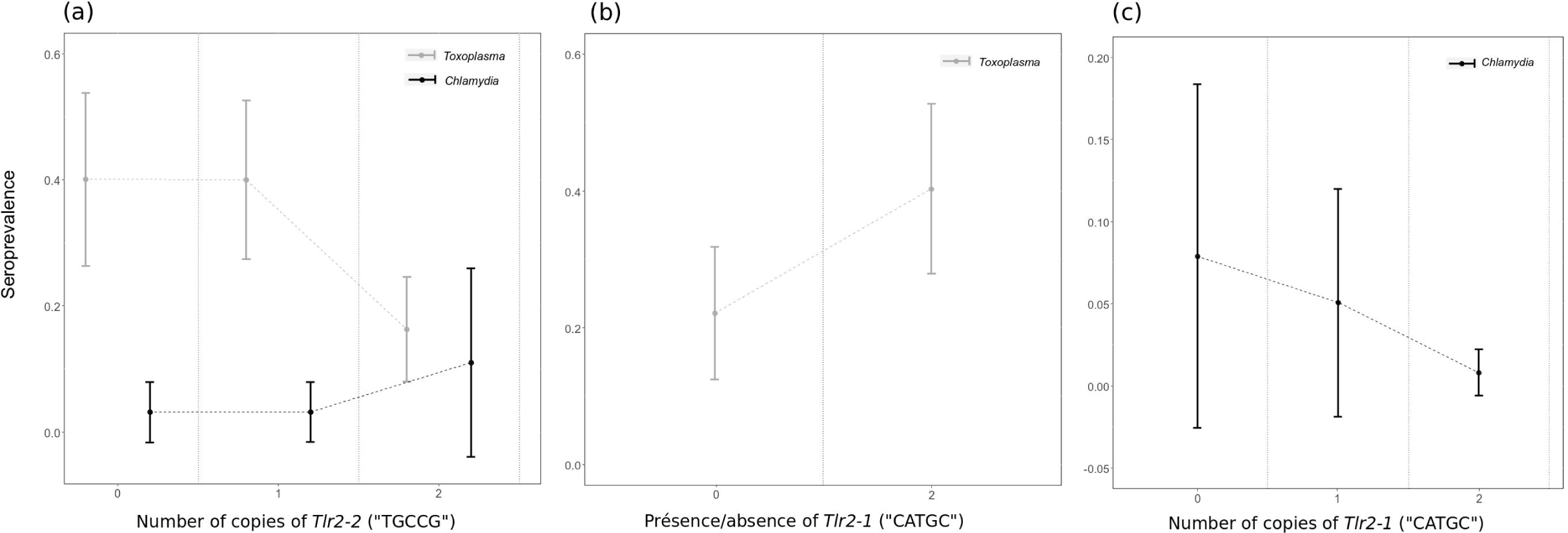
Genetic polymorphisms at *Tlr2* are associated with both *Chlamydia* and *Toxoplasma* serological status. (a) Predicted seroprevalence of *Toxoplasma* and *Chlamydia* in roe deer with the number of copies (0/1/2) of *Tlr2-2*, (b) Predicted seroprevalence of *Chlamydia* with the number of copies (0/1/2) of *Tlr2-1* (c) Predicted prevalence of *Toxoplasma* with the presence/absence of *Tlr2-1*.

Similarly to *Toxoplasma*, we found a strong relationship between *Tlr2* genotype and *Chlamydia* seroprevalence (Table 2) with the best (ΔAICc = −7.35 with non-genetic model) and second-best fitted models (ΔAICc = −5.96) including the number of copies of the “TGCCG” (*Tlr2-2*) and “CATCG” (*Tlr2-1*) haplotypes respectively. However, we observed the opposite pattern of association than for *Toxoplasma*: homozygous individuals for the most-common haplotype (*Tlr2-2*) had an increased probability to be seropositive (OR = 3.81 [1.13-12.89]) while individuals carrying *Tlr2-1* were less likely to be seropositive for *Chlamydia* (OR = 0.09 [0.01, 0.82] for “CATCG” homozygous deer) (Figure 1a, 1c, Table S8). The best-fitted model also revealed an association between *Tlr5* genotype and *Chlamydia* seroprevalence: roe deer with the “ACCTG” (*Tlr5-2*, 31%) haplotype were less likely to be Chlamydia seropositive (OR = 0.28[0.11, 0.72]). No association was observed when considering *Mhc-Drb* and *Tlr4* genotypes (neither number of alleles nor heterozygosity status).

## 4. DISCUSSION

Being evolutionary ancient and under strong functional constraints, genes involved in innate immune responses are expected to evolve primarily under purifying selection and to exhibit limited polymorphism (Parham, 2003). Yet, it is now quite clear that some of these genes, including *Tlr* genes encoding surface recognition receptors, may have relaxed selective constraints and show large nucleotide diversity, sometimes comparable to *Mhc* genes (Seabury *et al*., 2011). In a previous work, we have shown that the high sequence and allelic diversity observed at roe deer *Tlr* genes could be maintained through balancing selection (Quéméré *et al*., 2015). In this study, we developed a multi-gene/multi-pathogen association approach to bring new insights into the mechanisms underlying such balancing selection. Our results suggested a key role of directional selection on specific *Tlr* haplotypes rather than heterozygote advantage. We showed that different *Tlr* genes may be associated with resistance to the same microparasite (here *Tlr2* and *Tlr5* with *Chlamydia*) in agreement with the general idea that pathogens interact with many immune recognition receptors. We also revealed that different pathogens may act antagonistically on the polymorphism of a given gene (here *Tlr2* with *Toxoplasma* and *Chlamydia*).

### 4.1. High temporal and spatial variation in pathogen exposure

Our results revealed a high temporal heterogeneity in both pathogen seroprevalence at individual and population levels. In contrast with the traditional view of lifelong persistence of *Toxoplasma* infection (Tenter *et al*., 2000), we observed low individual repeatability of *Toxoplasma* seroprevalence at the within-individual level (R=0.21). This is in line with a previous study on the same population that reported frequent seroconversion with initially seropositive individuals becoming seronegative within 1 to 3 years (Sevila *et al*., 2014). *Toxoplasma* seroprevalence increased with roe deer age as frequently observed in wild ungulate populations, but it is most likely due to repeated exposure from the same environment rather than antibody persistence (Opsteegh *et al*., 2011; Gotteland *et al*., 2014). Overall, this suggests that the presence of antibodies in the study population would reflect relatively recent rather than long-term infection. The observed annual variation in pathogen prevalence at population level may be related to both climate and host factors. Meteorological conditions such as temperature or humidity are known to affect pathogen survival in the environment and may influence host transmission *via* the quantity of infected aerosols for *Chlamydia* (Tang, 2009) or oocysts for *Toxoplasma* (Gotteland *et al*., 2014). For example, Sevila et al. (2014) showed that *Toxoplasma* prevalence in roe deer was higher in mild and wet years and that *Chlamydia* prevalence increased in cold years. The high turnover of individuals in the population in relation with a strong harvest pressure may also partly explain why we observed a so high inter-annual variability in multi-cohort samples (Candela *et al*., 2014). Moreover, we also observed a strong variation in *Toxoplasma* exposure across sampling sectors that most likely results from landscape heterogeneity (e.g. exposure increase with the proportion of human dwellings within home range as a proxy of the cat presence, Sevila et al., 2014).

### 4.2 Predominant role of directional selection on *Tlr* gene variation

We revealed that *Tlr2* gene polymorphism is partly shaped by directional selection exerted by both *Toxoplasma* and *Chlamydia*. Roe deer carrying the most frequent *Tlr2* haplotype were less likely to be infected by *Toxoplasma* but more likely infected by *Chlamydia*. The opposite pattern was revealed for the second most frequent haplotype. These two abortive pathogens may have significant impact on the annual reproductive success of ungulates (Pioz *et al*., 2008). TLRs have been shown to activate the complement system, a component of innate immunity known to be important for resistance to microparasites during early infection (Raby *et al*., 2011). A higher affinity of a *Tlr* haplotype to *Toxoplasma* or *Chlamydia* ligands may result in a stronger complement response and thus in an increased resistance to these pathogens (Tschirren et al., 2013).

Spatial and temporal variations in pathogen-mediated directional selection suggest that the role of *Tlr2* haplotypes in roe deer resistance or susceptibility to *Toxoplasma* and *Chlamydia* may vary among years and among landscape structures where females established their home range. This supports the general idea that different habitats in terms of biotic and abiotic factors are likely to support distinct pathogen communities and so distinct pathogen-mediated-selection regime (Hedrick, 2002; Eizaguirre and Lenz, 2010). However, this result alone is not sufficient to demonstrate that fluctuating selection could favours the maintenance of *Tlr* genes’ balanced polymorphism in this population. Indeed, further data using longer time series are required to explicitly test whether *Toxoplasma* and *Chlamydia* prevalence are negatively correlated in space and/or time, which is a prerequisite for demonstrating that selection pressures are fluctuating (Spurgin and Richardson, 2010). Still few empirical studies have succeeded in providing evidence that fluctuating selection mediated immunogenetic variation, although they highlighted *Mhc* genetic diversity variation across space in a mosaic of habitats likely (Landry and Bernatchez, 2001; Alcaide *et al*., 2008; Eizaguirre and Lenz, 2010).

### 4.3 Antagonistic pleiotropy and potential immunogenetic trade-off

Interestingly, we observed that different haplotypes of a same *Tlr* gene (here *Tlr2*) may be associated with the infection status of the two pathogens. Pleiotropic effects in *Tlr2* were expected since this receptor is known to be implicated in the recognition of many bacterial, fungal, protozoa (de Oliviera Nascimento *et al*., 2012) including *Toxoplasma* (Yarovinsky, 2008) and *Chlamydia* (Darville *et al*., 2003; Beckett *et al*., 2012). Here, the most common *Tlr2* haplotype was associated with decreased *Toxoplasma* infection and increased *Chlamydia* infection while the concurrent haplotype showed the opposite trend. This result provides an example of antagonistic pleiotropy, which has been proposed as a widespread mechanism favouring the maintenance of genetic variation within populations (Rose, 1982; Roff and Fairbairn, 2007), in particular at immunity-related genes (Aderem and Ulevitch, 2000; Carter and Nguyen, 2011). Antagonistic pleiotropy has rarely been evidenced in natural populations (Turner *et al*., 2011) and, to our knowledge, this study is the first demonstration with regard to a *Tlr* gene. Two non-exclusive mechanisms may be invoked to explain this antagonistic effect: decrease of intracellular pathogen competition (Loiseau *et al*., 2008) and host immunogenetic trade-off (Kamath *et al*., 2014). *Toxoplasma* and *Chlamydia* use similar strategies to invade cells route and compete for the same nutrient pools in co-infected cells (Romano and Coppens, 2013). Therefore, antagonistic effects might arise because *Tlr2* gene alters the competitive interactions between the two pathogens. In other words, because individuals carrying the most frequent *Tlr2* haplotype are less infected by *Toxoplasma*, this may indirectly promote infection by *Chlamydia* without a direct role of the second most frequent haplotype (and *vice versa*). In a study of house sparrows, Loiseau et al. (2008) found antagonistic effects of a *Mhc* class I gene on multiple malarial parasite strains. In this particular case, deleterious ‘susceptibility alleles’ could be maintained in the population due to within-host competition between malaria parasite strains. Here, because we observed relatively few co-infections, it is unlikely that antagonistic effect may result from the competitive interactions between *Toxoplasma* and *Chlamydia*.

The most likely hypothesis is the presence of a host immunogenetic trade-off (Kamath *et al*., 2014). In this case, pathogens do not directly compete but *TLr2* genotype has a direct effect on the resistance/susceptibility to both pathogens. The allele inferred to be beneficial for decreasing *Toxoplasma* infection is also associated with increased susceptibility to *Chlamydia* (and *vice versa*). This scenario is reinforced by the fact that the two concurrent *Tlr2* haplotypes exhibit very distinct DNA and amino-acid sequences, what suggests marked functional differences. Similar patterns and processes have previously been described for ticks and nematodes infections (Kamath et al., 2014; Turner et al., 2011). To our knowledge, our study is the second one showing evidence of antagonistic selection at non-*Mhc* gene in a wild population (see for the other one Turner et al., 2011).

### 4.4 Limited role of other genes and selective mechanisms

While many empirical studies in a variety of organisms revealed signatures of pathogen-mediated directional or balancing in *Mhc* loci (see Bernatchez & Landry, 2003; Sommer, 2005; Spurgin & Richardson, 2010 for reviews), we did not find any association between *Mhc-Drb* and *Toxoplasma* or *Chlamydia*. This is congruent with the findings of Quéméré et al. (2015), who suggests that genetic drift and migration would the predominant contemporary forces shaping *Mhc* variation in roe deer with no signature of contemporary selection (in contrast to *Tlr* genes). Moreover, the few studies showing associations between *Mhc* class II genetic diversity and pathogen infection in other wild ungulates focused on strongyle parasites (Paterson *et al*., 1998; Charbonnel and Pemberton, 2005; Kamath *et al*., 2014) while we looked at interactions with micro-pathogens. This supports the general idea that MHC class II molecules principally bind exogenous antigens and are primarily involved in the immune response to extracellular pathogens (Hughes and Yeager, 1998).

Our data provided a weak support for the “heterozygous advantage hypothesis”. We noted that most authors revealing heterozygous advantage in immunity-related genes (most often on *Mhc* genes) studied animals co-infected with multiple pathogens (Hughes & Nei, 1992; McClelland et al., 2003; Froeschke & Sommer, 2012 but see Oliver et al., 2009). In our case, the lack of association with roe deer heterozygosity status could be due to the low individual’s probability of being exposed to both pathogens *Toxoplasma* and *Chlamydia* across their life-time, explaining the low fitness benefit of heterozygous genotypes. A limitation of this work is that we could not investigate the importance of other mechanisms such as negative frequency-dependent selection, because of insufficient statistical power to test the role of rare variants and the lack of long time series (see Charbonnel et al., 2005 for an example).

## 5. Conclusion

In conclusion, our study illustrates the importance of looking beyond *Mhc* genes (Acevedo-Whitehouse and Cunningham, 2006) and single snapshot gene/pathogen association (Maizels and Nussey, 2013) in wildlife immunogenetic studies. In the present case, we did not find any influence of *Mhc* diversity on the resistance to two microparasites. Moreover, we highlighted the importance of antagonistic effects on the maintenance of innate immunogenetic diversity, what could not have been possible using a classical single gene-single pathogen approach.

## ACKNOWLEDGMENTS

This work was supported by the French National Institute for Agricultural Research (INRA). We thank the local hunting associations, the Fédération Départementale des Chasseurs de Haute-Garonne. We thank numerous co-workers and volunteers for their assistance in field data collection, in particular J-M. Angibault, B. Cargnelutti, J-L. Rames, J. Joachim, B. Lourtet, D. Picot and N. Cebe. We also thank Karine Laroucau and Rachid Aaziz from ANSES the laboratory for animal health in Maison-Alfort (France) for their support and expertise on *C. abortus* results. We also thank Monica G Candela, Julie Sevila and co-workers for their support on laboratory work (serology). Genetic data used in this work were produced through molecular genetic analysis technical facilities of the labex “Centre Méditerranéen de l’Environnement et de la Biodiversité”.

## Data Archiving

Haplotype DNA sequences: Primers and Genbank accessions are in Table S1. SNP positions and characteristics are in Table S2. Genotypes of all individuals at each *Tlr* and *Mhc-Drb* gene and environmental/parasitological data will be shared on Dryad: Dryad entry XXXXXXXXXXX.

## Supporting information

**Table S1**. Summary of the 13 *Tlr* SNPs genotyped using the KASPar SNP genotyping system.

**Table S2**. Details of primer sequences, product size and Genbank accessions.

**Table S3**. Sample size per year and site for *Toxoplasma* and *Chlamydia*.

**Table S4**. Full list and performance of models tested for genetic association with *Chlamydia* and *Toxoplasma* seroprevalence.

**Table S5**. Prevalence of Toxoplasma and Chlamydia in the three sectors.

**Table S6**. Full list and performance of non-genetic models of *Chlamydia* and *Toxoplasma* seroprevalence.

**Table S7**. Parameter estimates from the best-fitted generalized linear mixed-effect describing variation in *Toxoplasma* seroprevalence as a function of age and number of copies of the *Tlr2-2* haplotype.

**Table S8**. Parameter estimates from the best-fitted generalized linear mixed-effect describing variation in *Chlamydia* seroprevalence as a function number of copies of the *Tlr2-2* and *Tlr5-2* haplotypes.

## REFERENCES

Acevedo-Whitehouse K, Cunningham A (2006). Is MHC enough for understanding wildlife immunogenetics? Trends Ecol Evol 21: 433–438.

Aderem A, Ulevitch RJ (2000). Toll-like receptors in the induction of the innate immune response. Nature 406: 782.

Akira S, Takeda K, Kaisho T (2001). Toll-like receptors: critical proteins linking innate and acquired immunity. Nat Immunol 2: 675–680.

Alcaide M, Edwards SV, Negro JJ, Serrano D, Tella JL (2008). Extensive polymorphism and geographical variation at a positively selected MHC class II B gene of the lesser kestrel (Falco naumanni). Mol Ecol 17: 2652–2665.

Antonides J, Mathur S, Sundaram M, Ricklefs R, DeWoody JA (2019). Immunogenetic response of the bananaquit in the face of malarial parasites. BMC Evol Biol 19: 107.

Barton K (2009). MuMIn: multi-model inference. R package version 1. 0. 0. Httpr-Forge R-Proj Orgprojectsmumin.

Bates D, Maechler M, Bolker B (2012). lme4: Linear mixed-effects models using S4 classes.

Beckett EL, Phipps S, Starkey MR, Horvat JC, Beagley KW, Foster PS, et al. (2012). TLR2, but not TLR4, is required for effective host defence against Chlamydia respiratory tract infection in early life. PLoS One 7: e39460.

Bernatchez L, Landry C (2003). MHC studies in nonmodel vertebrates: what have we learned about natural selection in 15 years? J Evol Biol 16: 363–377.

Brambilla A, Keller L, Bassano B, Grossen C (2018). Heterozygosity–fitness correlation at the major histocompatibility complex despite low variation in Alpine ibex (Capra ibex). Evol Appl 11: 631–644.

Candela MG, Serrano E, Sevila J, León L, Caro MR, Verheyden H (2014). Pathogens of zoonotic and biological importance in roe deer (Capreolus capreolus): Seroprevalence in an agro-system population in France. Res Vet Sci 96: 254–259.

Carter AJ, Nguyen AQ (2011). Antagonistic pleiotropy as a widespread mechanism for the maintenance of polymorphic disease alleles. BMC Med Genet 12: 160.

Charbonnel N, Pemberton J (2005). A long-term genetic survey of an ungulate population reveals balancing selection acting on MHC through spatial and temporal fluctuations in selection. Heredity 95: 377.

Coulon A, Guillot G, Cosson JF, Angibault JMA, Aulagnier S, Cargnelutti B, et al. (2006). Genetic structure is influenced by landscape features: empirical evidence from a roe deer population. Mol Ecol 15: 1669–1679.

Darville T, O’Neill JM, Andrews CW, Nagarajan UM, Stahl L, Ojcius DM (2003). Toll-like receptor-2, but not Toll-like receptor-4, is essential for development of oviduct pathology in chlamydial genital tract infection. J Immunol 171: 6187–6197.

Doherty PC, Zinkernagel RM (1975). Enhanced immunological surveillance in mice heterozygous at the H-2 gene complex. Nature 256: 50.

Dubey JP (2009). Toxoplasmosis in sheep—the last 20 years. Vet Parasitol 163: 1–14.

Dubois A, Galan M, Cosson J-F, Gauffre B, Henttonen H, Niemimaa J, et al. (2017). Microevolution of bank voles (Myodes glareolus) at neutral and immune-related genes during multiannual dynamic cycles: Consequences for Puumala hantavirus epidemiology. Infect Genet Evol 49: 318–329.

Eizaguirre C, Lenz TL (2010). Major histocompatibility complex polymorphism: dynamics and consequences of parasite-mediated local adaptation in fishes. J Fish Biol 77: 2023–2047.

Eizaguirre C, Lenz TL, Kalbe M, Milinski M (2012). Rapid and adaptive evolution of MHC genes under parasite selection in experimental vertebrate populations. Nat Commun 3: 621.

Froeschke G, Sommer S (2012). Insights into the complex associations between MHC class II DRB polymorphism and multiple gastrointestinal parasite infestations in the striped mouse. PLoS One 7: e31820.

Gazzinelli RT, Denkers EY (2006). Protozoan encounters with Toll-like receptor signalling pathways: implications for host parasitism. Nat Rev Immunol 6: 895–906.

Gervais L, Perrier C, Bernard M, Merlet J, Pemberton JM, Pujol B, et al. (2019). RAD-sequencing for estimating genomic relatedness matrix-based heritability in the wild: A case study in roe deer. Mol Ecol Resour.

Gotteland C, Aubert D, Gibert P, Moinet M, Klein F, Game Y, et al. (2014). Toxoplasmosis in natural populations of ungulates in France: prevalence and spatiotemporal variations. Vector-Borne Zoonotic Dis 14: 403–413.

Grueber CE, Wallis GP, Jamieson IG (2013). Genetic drift outweighs natural selection at toll-like receptor (TLR) immunity loci in a re-introduced population of a threatened species. Mol Ecol 22: 4470–4482.

Hedrick PW (1999). Antagonistic pleiotropy and genetic polymorphism: a perspective. Heredity 82: 126–133.

Hedrick PW (2002). Pathogen resistance and genetic variation at MHC loci. Evolution 56: 1902–1908.

Hedrick PW (2006). Genetic polymorphism in heterogeneous environments: the age of genomics. Annu Rev Ecol Evol Syst 37: 67–93.

Hedrick PW (2012). What is the evidence for heterozygote advantage selection? Trends Ecol Evol 27: 698–704.

Hewison A, Morellet N, Verheyden H, Daufresne T, Angibault J, Cargnelutti B, et al. (2009). Landscape fragmentation influences winter body mass of roe deer. Ecography 32: 1062–1070.

Hill AV (1991). HLA associations with malaria in Africa: some implications for MHC evolution. In: Molecular evolution of the major histocompatibility complex, Springer, pp 403–420.

Hughes AL, Nei M (1992). Models of host-parasite interaction and MHC polymorphism. Genetics 132: 863.

Hughes AL, Yeager M (1998). Natural selection and the evolutionary history of major histocompatibility complex loci. ront Biosci J Virtual Libr 3: d509–516.

Jepson A, Banya W, Sisay-Joof F, Hassan-King M, Nunes C, Bennett S, et al. (1997). Quantification of the relative contribution of major histocompatibility complex (MHC) and non-MHC genes to human immune responses to foreign antigens. Infect Immun 65: 872–876.

Johnston SE, Gratten J, Berenos C, Pilkington JG, Clutton-Brock TH, Pemberton JM, et al. (2013). Life history trade-offs at a single locus maintain sexually selected genetic variation. Nature 502: 93.

Kaisho T, Akira S (2004). Pleiotropic function of Toll-like receptors. Microbes Infect 6: 1388–1394.

Kamath PL, Turner WC, Küsters M, Getz WM (2014). Parasite-mediated selection drives an immunogenetic trade-off in plains zebras (Equus quagga). Proc R Soc Lond B Biol Sci 281: 20140077.

Klein SL (2000). The effects of hormones on sex differences in infection: from genes to behavior. Neurosci Biobehav Rev 24: 627–638.

Klein J, Figueroa F (1986). Evolution of the major histocompatibility complex. Crit Rev Immunol 6: 295–386.

Kloch A, Wenzel MA, Laetsch DR, Michalski O, Welc-Faleciak R, Piertney SB (2018). Signatures of balancing selection in toll-like receptor (TLRs) genes–novel insights from a free-living rodent. Sci Rep 8: 8361.

Kubinak JL, Ruff JS, Hyzer CW, Slev PR, Potts WK (2012). Experimental viral evolution to specific host MHC genotypes reveals fitness and virulence trade-offs in alternative MHC types. Proc Natl Acad Sci 109: 3422–3427.

Lam-Yuk-Tseung S, Gros P (2003). Genetic control of susceptibility to bacterial infections in mouse models. Cell Microbiol 5: 299–313.

Landry C, Bernatchez L (2001). Comparative analysis of population structure across environments and geographical scales at major histocompatibility complex and microsatellite loci in Atlantic salmon (Salmo salar). Mol Ecol 10: 2525–2539.

Levene H (1953). Genetic equilibrium when more than one ecological niche is available. Am Nat 87: 331–333.

Lewontin RC, Hubby JL (1966). A molecular approach to the study of genic heterozygosity in natural populations. II. Amount of variation and degree of heterozygosity in natural populations of Drosophila pseudoobscura. Genetics 54: 595.

Loiseau C, Zoorob R, Garnier S, Birard J, Federici P, Julliard R, et al. (2008). Antagonistic effects of a Mhc class I allele on malaria-infected house sparrows. Ecol Lett 11: 258–265.

Maizels RM, Nussey DH (2013). Into the wild: digging at immunology’s evolutionary roots. Nat Immunol 14: 879.

McClelland EE, Penn DJ, Potts WK (2003). Major histocompatibility complex heterozygote superiority during coinfection. Infect Immun 71: 2079–2086.

Miller CM, Boulter NR, Ikin RJ, Smith NC (2009). The immunobiology of the innate response to Toxoplasma gondii. Int J Parasitol 39: 23–39.

Møller AP, Saino N (2004). Immune response and survival. Oikos 104: 299–304.

Moore AJ, Moore PJ (1999). Balancing sexual selection through opposing mate choice and male competition. Proc R Soc Lond B Biol Sci 266: 711–716.

Morellet N, Bonenfant C, Börger L, Ossi F, Cagnacci F, Heurich M, et al. (2013). Seasonality, weather and climate affect home range size in roe deer across a wide latitudinal gradient within E urope. J Anim Ecol 82: 1326–1339.

Mun H-S, Aosai F, Norose K, Chen M, Piao L-X, Takeuchi O, et al. (2003). TLR2 as an essential molecule for protective immunity against Toxoplasma gondii infection. Int Immunol 15: 1081–1087.

Netea M, Wijmenga C, O’Neill L (2012). Genetic variation in Toll-like receptors and disease susceptibility. Nat Immunol 13: 535–542.

Oliver MK, Telfer S, Piertney SB (2009). Major histocompatibility complex (MHC) heterozygote superiority to natural multi-parasite infections in the water vole (Arvicola terrestris). Proc R Soc Lond B Biol Sci 276: 1119–1128.

de Oliviera Nascimento L, Massari P, Wetzler LM (2012). The role of TLR2 in infection and immunity. Front Immunol 3: 79.

Opsteegh M, Swart A, Fonville M, Dekkers L, Van Der Giessen J (2011). Age-related Toxoplasma gondii seroprevalence in Dutch wild boar inconsistent with lifelong persistence of antibodies. PLoS One 6: e16240.

Parham P (2003). Innate immunity: the unsung heroes. Nature 423: 20–20.

Paterson S, Wilson K, Pemberton JM (1998). Major histocompatibility complex variation associated with juvenile survival and parasite resistance in a large unmanaged ungulate population (Ovis aries L.). Proc Natl Acad Sci 95: 3714–3719.

Pedersen AB, Greives TJ (2008). The interaction of parasites and resources cause crashes in a wild mouse population. J Anim Ecol 77: 370–377.

Phillips KP, Cable J, Mohammed RS, Herdegen-Radwan M, Raubic J, Przesmycka KJ, et al. (2018). Immunogenetic novelty confers a selective advantage in host–pathogen coevolution. Proc Natl Acad Sci: 201708597.

Pioz M, Loison A, Gauthier D, Gibert P, Jullien J-M, Artois M, et al. (2008). Diseases and reproductive success in a wild mammal: example in the alpine chamois. Oecologia 155: 691–704.

Piry S, Guivier E, Realini A, Martin J (2012). SESAME Barcode: NGS-oriented software for amplicon characterization– application to species and environmental barcoding. Mol Ecol Resour 12: 1151–1157.

Quéméré E, Galan M, Cosson J-F, Klein F, Aulagnier S, Gilot-Fromont E, et al. (2015). Immunogenetic heterogeneity in a widespread ungulate: the European roe deer (Capreolus capreolus). Mol Ecol.

Raby A-C, Holst B, Davies J, Colmont C, Laumonnier Y, Coles B, et al. (2011). TLR activation enhances C5a-induced pro-inflammatory responses by negatively modulating the second C5a receptor, C5L2. Eur J Immunol 41: 2741–2752.

Roff DA, Fairbairn DJ (2007). The evolution of trade-offs: where are we? J Evol Biol 20: 433–447.

Romano JD, Coppens I (2013). Host Organelle Hijackers: a similar modus operandi for Toxoplasma gondii and Chlamydia trachomatis: co-infection model as a tool to investigate pathogenesis. Pathog Dis 69: 72–86.

Rose MR (1982). Antagonistic pleiotropy, dominance, and genetic variation. Heredity 48: 63.

Seabury CM, Bhattarai EK, Taylor JF, Viswanathan GG, Cooper SM, Davis DS, et al. (2011). Genome-wide polymorphism and comparative analyses in the white-tailed deer (Odocoileus virginianus): a model for conservation genomics. Plos One 6: e15811.

Sevila J, Richomme C, Hoste H, Candela MG, Gilot-Fromont E, Rodolakis A, et al. (2014). Does land use within the home range drive the exposure of roe deer (Capreolus capreolus) to two abortive pathogens in a rural agro-ecosystem? Acta Theriol (Warsz) 59: 571–581.

Sin YW, Annavi G, Dugdale HL, Newman C, Burke T, MacDonald DW (2014). Pathogen burden, co-infection and major histocompatibility complex variability in the European badger (Meles meles). Mol Ecol 23: 5072–5088.

Sommer S (2005). The importance of immune gene variability (MHC) in evolutionary ecology and conservation. Front Zool 2: 16.

Spurgin L, Richardson D (2010). How pathogens drive genetic diversity: MHC, mechanisms and misunderstandings. Proc Biol Sci 277: 979–988.

Stoffel MA, Nakagawa S, Schielzeth H (2017). rptR: Repeatability estimation and variance decomposition by generalized linear mixed-effects models. Methods Ecol Evol 8: 1639–1644.

Takahata N, Nei M (1990). Allelic genealogy under overdominant and frequency-dependent selection and polymorphism of major histocompatibility complex loci. Genetics 124: 967–978.

Tang JW (2009). The effect of environmental parameters on the survival of airborne infectious agents. J R Soc Interface 6: S737–S746.

Tenter AM, Heckeroth AR, Weiss LM (2000). Toxoplasma gondii: from animals to humans. Int J Parasitol 30: 1217–1258.

Tollenaere C, Bryja J, Galan M, Cadet P, Deter J, Chaval Y, et al. (2008). Multiple parasites mediate balancing selection at two MHC class II genes in the fossorial water vole: insights from multivariate analyses and population genetics. J Evol Biol 21: 1307–1320.

Trowsdale J, Parham P (2004). Mini-review: defense strategies and immunity-related genes. Eur J Immunol 34: 7–17.

Tschirren B, Andersson M, Scherman K, Westerdahl H, Mittl P, Raberg L (2013). Polymorphisms at the innate immune receptor TLR2 are associated with Borrelia infection in a wild rodent population. Proc Biol Sci 280: 20130364.

Turner AK, Begon M, Jackson JA, Bradley JE, Paterson S (2011). Genetic Diversity in Cytokines Associated with Immune Variation and Resistance to Multiple Pathogens in a Natural Rodent Population. PLoS Genet 7: e1002343.

Wegner KM, Kalbe M, Kurtz J, Reusch TB, Milinski M (2003). Parasite selection for immunogenetic optimality. Science 301: 1343–1343.

Wellenreuther M (2017). Balancing selection maintains cryptic colour morphs. Mol Ecol 26: 6185–6188.

Yang X, Joyee A (2008). Role of toll-like receptors in immune responses to chlamydial infections. Curr Pharm Des 14: 593–600.

Yarovinsky F (2008). Toll-like receptors and their role in host resistance to Toxoplasma gondii. Immunol Lett 119: 17–21.

Zuur A, Ieno EN, Walker N, Saveliev AA, Smith GM (2009). Mixed effects models and extensions in ecology with R. Springer Science & Business Media.

