## Supplementary material for "Antagonistic pathogen-mediated selection favours the maintenance of innate immune gene polymorphism in a widespread wild ungulate": TableS1-S2-S3-S5-S6-S7-S8

**Supporting information**

Table S1. Summary of 13 *Tlr* SNPs genotyped using the KASPar SNP genotyping system.

| Gene | SNP^1^ | AA site^2^ | Codon Change | Syn/Nonsyn^3^ |
| --- | --- | --- | --- | --- |
| TLR2 | TLR2 238 C/T | 113 | TCT(Ser) > CCT(Pro) | Non Syn |
|  | TLR2 1540 A/G | 547 | GGG(Gly) > AGG(Arg) | Non Syn |
|  | TLR2 1366 C/T | 489 | CTG(Leu) > TTG(Leu) | Syn |
|  | TLR2 1533 C/T | 544 | TTC(Phe) > TTT(Phe) | Syn |
|  | TLR2 1637 A/G | 579 | CGG(Arg) > CAG(Glu) | Non Syn |
| TLR4 | TLR4 46 C/G | 128 | GAG (Glu) > CAG (Gln) | Non Syn |
|  | TLR4 768 A/G | 368 | GAA(Glu) > CAG(Glu) | Syn |
|  | TLR4 1489 A/G | 608 | GCA(Ala) > ACA(Thr) | Non Syn |
| TLR5 | TLR5 280 A/G | 122 | ACT(Thr) > GCT(Ala) | Non Syn |
|  | TLR5 415 C/T | 167 | CGG(Arg)>TGG(Trp) | Non Syn |
|  | TLR5 1239 C/T | 441 | AAC(Asn)>AAT(Asn) | Syn |
|  | TLR5 1869 C/G | 651 | ACC(Thr)> ACT(Thr) | Syn |
|  | TLR5 2350 A/G | 812 | GTC (Val) > ATC (Ile) | Non Syn |

^1^ SNP position relative to start of the roe deer sequence produced with primers in Table S2. ^2^ Position of SNP codon relative to aligned mouse amino acid sequence. ^3^ Denotes whether the SNP is synonymous or non-synonymous.

Table S2. Details of primer sequences, product size and genbank accessions.

| Gene | Primer name | Primer sequence (5’-3’) | Primer position | Product size (pb) | GENBANK Accession |
| --- | --- | --- | --- | --- | --- |
| *Tlr2* | TLR2a-F  TLR2a-R  TLR2b-F  TLR2b-R  TLR2c-F  TLR2c-R | CATCAGCCTGTCCACRGAAG  GTGACTCTTCTAACTCTGCCTG  GGAGACGTTGACAATACGGA  CTTGCCAGGAACGAAGTCTC  TTTGCTCCTGTGACTTCCTG  GCCACTCCAGGTAGGTCYTG | exon 1  exon 1  exon 1  exon 1  exon 1  exon 1 | 987  1123  687 | KM488227-> KM488232 |
| *Tlr3* | TLR3a-F  TLR3a-R  TLR3b-F  TLR3b-R | GGCCTCTCTCTGAACAATGC  GCCGAGCTAAGTTGTTATGC  TTCCGAACCTGGTCATTCTG  TGCTCCTTCTGATGCTGTTC | exon 4  exon 4  exon 4  exon 4 | 961  915 | KM488233 -> KM488235 |
| *Tlr4* | TLR4a-F2  TLR4a-R  TLR4b-F  TLR4b-R  TLR4c-F  TLR4c-R2 | CCACCTCTCCACCTTGATAC  GAACTGAGCTTCAATGCAGG  GTGCAACCTGACCATTGAAC  TCCTGGCTCGAGTAGATGAC  GACCTCTCTAAGTGTCAACTGG  CTGCTGTTCCTTCTGGACTC | exon 3  exon 3  exon 3  exon 3  exon 3  exon 3 | 750  1215  982 | KM488236 -> KM488245 |
| *Tlr5* | TLR5a-F  TLR5a-R  TLR5b-F  TLR5b-R  TLR5c-F  TLR5c-R | CCTGACCAGTCCTGCACTTG  TGAGAACTTGGAGGTTGTCG  GACACTCCAGGAGCTGAAGG  AGGAACAGAGTCAAGGTGACAG  CATCTGTGAGTGTGAACTTAGTGC  GCAGCTGAATAGCACTGTCTT | exon 1  exon 1  exon 1  exon 1  exon 1  exon 1 | 986  1035  812 | KM488246 -> KM488256 |
| *Mhc-Drb* | LA31-Fm  LA32-Rm | GATCCTCTCTCTGCAGCACATTTCCT  TTCGCGTCACCTCGCCGCTG | exon 2  exon 2 | 296 | KM488213 -> KM488222 |

**Table S3a. Sample size per year and Sector for *Toxoplasma***

| Sector | 2008 | 2009 | 2010 | 2011 | 2012 | 2013 | 2014 | 2015 | 2016 | Total |
| --- | --- | --- | --- | --- | --- | --- | --- | --- | --- | --- |
| Closed | 10 | 10 | 13 | 13 | 5 | 7 | 9 | 14 | 4 | 85 |
| Open | 27 | 29 | 13 | 21 | 17 | 20 | 12 | 27 | 9 | 175 |
| Mixed | 3 | 3 | 17 | 16 | 20 | 11 | 10 | 8 | 3 | 91 |
| Total | 40 | 42 | 43 | 50 | 42 | 38 | 31 | 49 | 16 | **351** |

**Table S3b. Sample size per Year and Sector for *Chlamydia***

| Sector | 2008 | 2009 | 2010 | 2011 | 2012 | 2013 | Total |
| --- | --- | --- | --- | --- | --- | --- | --- |
| Closed | 10 | 10 | 13 | 13 | 5 | 0 | 51 |
| Open | 27 | 29 | 13 | 21 | 17 | 14 | 121 |
| Mixed | 3 | 2 | 17 | 16 | 20 | 2 | 60 |
| Total | 40 | 41 | 43 | 50 | 42 | 16 | **232** |

Table S5. Prevalence of *Toxoplasma* and *Chlamydia* in the three sectors.

| Site_ID | *Toxoplasma* prevalence | *Chlamydia* prevalence |
| --- | --- | --- |
| Closed | 0.29 | 0.15 |
| Open | 0.45 | 0.11 |
| Mixed | 0.27 | 0.20 |

**Table S6a. Full list and performance of non-genetic generalized linear mixed-effect models of *Toxoplasma* seroprevalence**. The selected models occur in bold. ‘k’ refers to the number of model parameters. The selected models occur in bold.

| Model | k | ΔAICc | AICcWt |
| --- | --- | --- | --- |
| age + year + sector | **13** | **0** | **0.565** |
| age + year + sector + sex | 14 | 2.21 | 0.187 |
| age + year | 11 | 3.23 | 0.113 |
| year+ sector | 12 | 5.2 | 0.042 |
| age + year | 12 | 5.51 | 0.036 |
| year + sector | 10 | 5.79 | 0.031 |
| year + sector + sex | 13 | 6.81 | 0.019 |
| year + sex | 11 | 8.74 | 0.007 |
| age + sector | 5 | 49.47 | 0 |
| sector | 4 | 50.41 | 0 |
| age + sector + sex | 6 | 51.53 | 0 |
| sector + sex | 5 | 52.46 | 0 |
| constant | 2 | 53.43 | 0 |
| age | 3 | 53.78 | 0 |
| sex | 3 | 55.42 | 0 |
| age + sector | 4 | 55.78 | 0 |

**Table S6b. Full list and performance of non-genetic generalized linear mixed-effect models of *Chlamydia* seroprevalence**. The selected models occur in bold. ‘k’ refers to the number of model parameters. The selected models occur in bold.

|  | k | ΔAICc | AICcWt |
| --- | --- | --- | --- |
| year | **6** | **0** | **0.44** |
| year + sex | 7 | 1.65 | 0.193 |
| age + year | 7 | 1.82 | 0.177 |
| age + year + sex | 8 | 3.5 | 0.076 |
| year + sector | 8 | 4.05 | 0.058 |
| year + sector + sex | 9 | 5.84 | 0.024 |
| age + year + sector | 9 | 5.93 | 0.023 |
| age + year + sector + sex | 10 | 7.75 | 0.009 |
| constant | 1 | 39.7 | 0 |
| sector | 3 | 40.77 | 0 |
| sex | 2 | 41.53 | 0 |
| age | 2 | 41.57 | 0 |
| age + sector | 4 | 42.48 | 0 |
| sector + sex | 4 | 42.74 | 0 |
| age + sex | 3 | 43.42 | 0 |
| age + sector + sex | 5 | 44.48 | 0 |

**Table S7.** Parameter estimates from the best glmer describing variation in *Toxoplasma* seroprevalence as a function of age and number of copies of the *Tlr2-2* haplotype.

|  | Coefficient | SE | Z-value | Pr(>\|z\|) |
| --- | --- | --- | --- | --- |
| Intercept | -0.4 | 0.57 | -0.71 | 0.48 |
| AGE: juvenile category | -0.92 | 0.38 | -2.41 | 0.016 |
| nb(Tlr2-2): 1 copy category | -0.0036 | 0.39 | -0.009 | 0.99 |
| nb(Tlr2-2): 2 copies category | -1.23 | 0.49 | -2.48 | 0.013 |

**Table S8**. Parameter estimates from the best glmer describing variation in *Chlamydia* seroprevalence as a function number of copies of the *Tlr2-2* and *Tlr5-2* haplotypes.

|  | Coefficient | SE | Z-value | Pr(>\|z\|) |
| --- | --- | --- | --- | --- |
| Intercept | -3.43 | 1.56 | -2.19 | 0.03 |
| nb(Tlr2-2): 1 copy category | 0.01 | 0.59 | 0.02 | 0.98 |
| nb(Tlr2-2): 2 copies category | 1.34 | 0.62 | 2.16 | 0.03 |
| nb(Tlr5-2): 1 copy category | -1.29 | 0.49 | -2.62 | 0.01 |
| nb(Tl5-2): 2 copies category | -0.13 | 0.99 | -0.14 | 0.89 |
